# Age and ketogenic diet have dissociable effects on synapse-related gene expression between hippocampal subregions

**DOI:** 10.1101/718445

**Authors:** Abbi R. Hernandez, Caesar M. Hernandez, Leah M. Truckenbrod, Keila T. Campos, Joseph A. McQuail, Jennifer L. Bizon, Sara N. Burke

## Abstract

As the number of individuals living beyond the age of 65 is rapidly increasing, so is the need to develop strategies to combat the age-related cognitive decline that may threaten independent living. Although the link between altered neuronal signaling and age-related cognitive impairments is not completely understood, it is evident that declining cognitive abilities are at least partially due to synaptic dysfunction. Aging is accompanied by well-documented changes in both excitatory and inhibitory synaptic signaling across species. Age-related synaptic alterations are not uniform across the brain, however, with different regions showing unique patterns of vulnerability in advanced age. In the hippocampus, increased activity within the CA3 subregion has been observed across species, and this can be reversed with anti-epileptic medication (Bakker et al., 2012). In contrast to CA3, the dentate gyrus shows reduced activity with age and declining metabolic activity. Ketogenic diets have been shown to decrease seizure incidence and severity in epilepsy, improve metabolic function in diabetes type II, and improve cognitive function in aged rats. This link between neuronal activity and metabolism suggests that metabolic interventions may be able to ameliorate synaptic signaling deficits accompanying advanced age. We therefore investigated the ability of a dietary regimen capable of inducing nutritional ketosis and improving cognition to alter synapse-related gene expression across the dentate gyrus, CA3 and CA1 subregions of the hippocampus. Following 12 weeks of a ketogenic or calorie-matched standard diet, RTq-PCR was used to quantify expression levels of excitatory and inhibitory synaptic signaling genes within CA1, CA3 and dentate gyrus. While there were no age or diet-related changes in CA1 gene expression, expression levels were significantly altered within CA3 by age and within the dentate gyrus by diet for several genes involved in presynaptic glutamate regulation and postsynaptic excitation and plasticity. These data demonstrate subregion-specific alterations in synaptic signaling with age and the potential for a ketogenic diet to alter these processes in dissociable ways across different brain structures that are uniquely vulnerable in older animals.

## Introduction

Although declining cognitive function is common among older adults, across species, significant neuron loss is not observed in most areas of the brain, including the hippocampus (Keuker et al., 2004; Merrill et al., 2001; Mohammed and Santer, 2001; Pannese, 2011; Rapp et al., 2002; Rapp and Gallagher, 1996). While cell number in the hippocampus remains stable throughout the lifespan, relative to younger animals, aged primates and rodents show well-documented changes in synaptic structure and function within this brain region that relate to behavioral impairments (Barnes, 2003; Morrison and Baxter, 2012; Pannese, 2011; Rosenzweig and Barnes, 2003). These findings point to hippocampal synapses as key players in age-related cognitive deficits, suggesting that therapies designed to improve synaptic function in this structure could mitigate age-related behavioral decline.

While the entire hippocampus undergoes alterations in synaptic structure, function and plasticity with advancing age, these changes are not uniform between the different subregions (for review, see Burke and Barnes, 2006; Rosenzweig and Barnes, 2003). Synaptophysin, a presynaptic marker, is significantly reduced in old compared to young rats in both CA3 and the dentate gyrus, but remains stable in aged CA1 (Smith et al., 2000). In the dentate gyrus, there is also a significant age-related decrease in synaptic number within the molecular layer (Geinisman et al., 1992) that correlates with impaired spatial memory (Geinisman et al., 1986). This reduction in dentate gyrus synapses is likely related to loss of perforant path fibers from the entorhinal cortex that terminate in the outer and middle molecular layers (Barnes and McNaughton, 1980; Foster et al., 1991; Yassa et al., 2010, 2011). In the CA3 subregion, in addition to the loss of synapses in older animals, pyramidal cells show higher firing rates (Thomé et al., 2016; Wilson, 2005). Moreover, there is an increase in intrinsic excitability (Simkin et al., 2015; Villanueva-Castillo et al., 2017), and interneurons lose their phenotypic expression profiles (Cadacio et al., 2003; Koh et al., 2014; Spiegel et al., 2013; Thomé et al., 2016). Although the Schaffer collateral to CA1 pyramidal cell synapse does not exhibit an age-related decrease in total number (Geinisman et al., 2004), perforated synapses have reduced postsynaptic densities that correlate with cognitive impairment (Nicholson et al., 2004). These morphological data and the associated age-related decrease in the excitatory post-synaptic field potential that is evoked from Schaffer collateral stimulation (Barnes et al., 1992; Landfield et al., 1986) suggest that a portion of CA1 synapses may become nonfunctional or silent in advanced age (Rosenzweig and Barnes, 2003).

At the level of gene expression, there is also evidence that advanced age leads to alterations that are hippocampal subregion dependent. Overall gene expression profiles obtained from microarray analysis of individual hippocampal subregions show that while many genes are differentially-regulated in old compared to young rats, a higher number of genes are altered with age in CA3 relative to CA1 and the dentate gyrus. Moreover, age-related changes in gene expression in CA3 account for more variability in spatial memory performance (Haberman et al., 2011, 2013). Interestingly, the pathways in which multiple genes are affected also differs across subregions (Haberman et al., 2011), indicating that different cell signaling properties are uniquely vulnerable in old age between CA1, CA3 and the dentate gyrus. Another study using next-generation RNA sequencing, however, identified the dentate as the subregion with the greatest number of age-related changes in gene expression (Ianov et al., 2016). While CA1 appears to have fewer age-associated changes in synaptic function and gene expression relative to the other subregions, there is evidence that metabolism and plasticity-related gene expression in this subregion is altered between young and older rats, and that these changes correlate with impaired cognitive performance (Blalock et al., 2003).

The importance of synaptic activity for behavior is highlighted by the brain’s disproportionate use of metabolic resources for this process. A wide body of literature supports age-related impairments in metabolism within the hippocampus (Gage et al., 1984; Goyal et al., 2017; Hernandez et al., 2018a; Rasgon et al., 2005). Because synaptic transmission is so metabolically costly (Attwell and Laughlin, 2001; Howarth et al., 2012), it is possible that age-related impairments in cerebral glucose metabolism contribute to age-related synaptic dysfunction. Although cerebral glucose metabolism declines with age, the ability to utilize ketone bodies for energy production does not (Castellano et al., 2015; Cunnane et al., 2016; Lying-Tunell et al., 1981; Ogawa et al., 1996). Therefore, the potential to bypass impaired glycolysis through the elevation of ketone body synthesis and promotion of fat oxidation may alter the expression of hippocampal synaptic signaling-related genes (Hernandez et al., 2018a) to ultimately improve cognitive function (Hernandez et al., 2018b).

Ketogenic diets are high in fat and low in carbohydrates, switching the body’s primary fuel source from glucose to ketone bodies. Ketogenic diets have been utilized to alter neuronal signaling patterns in patients with epilepsy for almost a century (Freeman et al., 1998). More recent evidence indicates that it can alter both excitatory and inhibitory signaling-related proteins within young and aged brains (Hernandez et al., 2018a, 2018b). These alterations in protein expression have been shown to be both age- and brain region-specific, with more pronounced effects in the hippocampus than the prefrontal cortex. Quantification of proteins, however, does not easily afford the ability to delineate between hippocampal subregions, as is possible by examining gene expression at the level of RNA. This distinction is critical as previous work has suggested that ketone bodies may differentially affect the CA1 and dentate gyrus subregions. Specifically, the addition of medium chain triglyceride oil to rodent chow, which moderately increases ketone body levels in aged rats, was reported to increase the number of synaptic contacts and improve mitochondrial functioning within the dentate gyrus, but reduce synaptic contacts in CA1 (Balietti et al., 2008). Relatedly, caloric restriction, which can induce fat oxidation and elevate ketone body levels, has also been shown to have a greater impact on gene expression in CA3 and the dentate gyrus compared to CA1 (Zeier et al., 2011). Thus, there is a clear link between synaptic function, age and metabolism that differs across hippocampal subregions. The extent to which switching neurometabolism from glycolysis to fat oxidation impacts the expression of genes related to synaptic signaling, however, has not been empirically examined in young and aged animals.

To investigate the role of a ketogenic dietary intervention on synaptic signaling-related gene expression within hippocampal subregions, young and aged rats were placed on an isocaloric ketogenic or control standard diet for 12 weeks. Excitatory and inhibitory synaptic signaling-related gene expression was quantified within the dentate gyrus, CA3 and CA1 regions of the hippocampus with RTq-PCR. While there were no significant age- or diet-dependent changes within CA1, there were several age-related changes in gene expression within CA3 and several diet-related changes observed within the dentate gyrus. These data add to a growing body of work establishing the CA3 hippocampal subregion as being particularly vulnerable to dysfunction with age and provides new insights into how promoting fat oxidation over glycolysis leads to dissociable effects between the different hippocampal subregions.

## Methods

### Subjects and handling

Young (4 months; n = 15) and aged (20 months: n = 16) male Fischer 344 × Brown Norway F1 Hybrid rats from the NIA colony at Charles River were used for the current study. One week after arrival to the colony, 7-8 rats per age group were placed on either the ketogenic or standard diet. Rats were housed individually and maintained on a 12-hr reverse light/dark cycle. Feeding and all data collection, including tissue harvesting, were conducted during the rats’ dark cycle. All experimental procedures were performed in accordance with National Institutes of Health guidelines and were approved by Institutional Animal Care and Use Committees at the University of Florida.

### Diet

Prior to diet administration, rats were randomly assigned to either a high-fat, low-carbohydrate ketogenic diet (KD; 75.85% fat, 20.12% protein, 3.85% carbohydrate; Lab Supply; 5722, Fort Worth, Texas) mixed with medium chain triglyceride (MCT) oil (Neobee 895, Stephan, Northfield, Illinois) or a calorically and micronutrient matched standard diet (SD; 16.35% fat, 18.76% protein, 64.89% carbohydrate; Lab Supply; 1810727, Fort Worth, Texas; for details on diet, see Hernandez et al., 2018). Rats were weighed daily and fed equivalent calories (~51 kcal/day) at the same time each day for 12 weeks. This caloric quantity, when fed once daily, is sufficient for healthy weight maintenance on both a KD and SD (Hernandez et al., 2018a). Importantly, since all age and diet groups were calorically matched, energy intake could not account for group differences in the data. Access to water was *ad libitum* and all groups consumed comparable amounts of water (Hernandez et al., 2018a).

### Confirmation of nutritional ketosis

To confirm rats were in nutritional ketosis, blood glucose and ketone body (β-hydroxybutyrate; BHB) levels were quantified on the day of sacrifice and tissue collection. For each measurement, one drop of blood was taken directly from the trunk immediately following decapitation for each test. Blood was collected directly onto the appropriate test strip (Abbott Diabetes Care, Inc, Alameda, CA; glucose SKU#: 9972865 and ketone SKU#: 7074565), which was placed into a Precision Xtra blood monitoring system (Abbott Diabetes Care, Inc, Alameda, CA; SKU#: 9881465).

### Tissue Collection

On the day of sacrifice, rats were placed into a bell jar containing isoflurane-saturated cotton (Abbott Laboratories, Chicago, IL, USA), separated from the animal by a wire mesh shield. Animals lost righting reflex within 30 seconds of being placed within the jar and were then immediately euthanized by rapid decapitation. The left and right hemispheres of the hippocampus were isolated bilaterally, immediately frozen on dry ice and stored at −80°C until use. The left hippocampus was left whole for Western blotting, and the right hippocampus was subdivided into hippocampal subregions CA1, CA3 and dentate gyrus for RNA quantification.

### Reverse Transcription and PCR Expression Assay

On the day of RNA extraction, tissue was transferred to QIAzol Lysis Reagent (PN: 79306, Qiagen, Frederick, MD, USA) and total RNA was isolated using the RNEasy Plus Mini kit according to the manufacturer’s protocol (PN: 73404, Qiagen). RNA concentration was determined with the use of a Quibit Fluorometer (ThermoFisher, Waltham, MA USA). The yield of RNA was consistent and reproduced across groups. The average RNA integrity number (RIN) was determined by TapeStation (Agilent Biosciences, Santa Clara, CA, USA) was 8.91 (±0.29 SD), and no sample had a RIN lower than 8.2.

From each sample, 600 ng of RNA was used to make cDNA using the RT^2^ First Strand cDNA Synthesis Kit (PN: 330411, Qiagen). Relative gene expression was measured using RT^2^ Profiler low-density PCR plates preloaded with qPCR primer assays for genes encoding GABA- and glutamate-related targets (PN: PARN-152ZA, Qiagen). Thermal cycling and data collection were accomplished using an ABI Real-Time PCR 7300.

### Gene Exclusion Criteria and Statistical Analyses

Only RT-qPCR plates that passed the PCR array reproducibility, reverse transcription efficiency, and genomic DNA contamination quality control parameters set by Qiagen’s methods (RT^2^ Profiler PCR Array Data Analysis v3.5), as well as those reactions that produced the predicted peak by melting temperature (T_m_) curve analysis were included in the final analyses. Genes were normalized to the house-keeping gene *RPLP1*. Any samples with cycle threshold (C_t_) values for *RPLP1* that were statistical outliers (as determined by the ROUT method with a false discovery rate (FDR) of q = 0.005) were excluded from downstream analyses. After necessary exclusions, the final group sizes across the different hippocampal subregions are shown in Table 1. Note, that for some genes the exclusion of outliers may have led to slightly smaller group sizes.

Each gene included in the RT-qPCR plates was cross-referenced with the Allen Brain Institute’s online *in situ* hybridization atlas (http://mouse.brain-map.org/) and those not expressed in hippocampus were used to set the lowest C_t_ considered detectable. As such, the final number of genes tested per region was 60 (DG), 58 (CA3), and 55 (CA1). After normalization with *RPLP1* and transformation using the 2^(-ΔCt)^ method, values were standardized and expressed as Z-scores. Two-factor ANOVAs (age × diet) were used to compare expression of gene transcripts between group samples in DG, CA3 and CA1 separately using the Benjamini-Hochberg method to correct for multiple comparisons with a FDR of q=0.005 (Benjamini and Hochberg, 1995; Storey and Tibshirani, 2003).

**Table 1:** Final group sizes for PCR gene analysis.

| Group | Young (n) |  | Aged (n) |  |
| --- | --- | --- | --- | --- |
| Diet | Control | Keto | Control | Keto |
| DG | 7 | 7 | 8 | 4 |
| CA3 | 7 | 6 | 8 | 6 |
| CA1 | 6 | 8 | 6 | 4 |

### Gene Functional Enrichment Analysis

The Database for Annotation, Visualization and Integrated Discovery (DAVID, v6.8) was used to determine any significantly enriched pathways associated with genes that met the FDR criteria (Huang et al., 2009a, 2009b). Specifically, DAVID enabled the identification of the most robustly enriched Kyoto Encyclopedia of Genes and Genomes (KEGG) pathway per gene set. To simplify downstream analyses and interpretations, only the top FDR-ranked pathways per gene list were considered. Significantly enriched KEGG pathways were validated using a second enrichment analysis tool such as Enrichr (Chen et al., 2013; Cowley Michael et al.; Kuleshov et al., 2016).

### Evaluation of ROCK Protein Levels

Whole hippocampal homogenates were used from 5 of these rats per age and diet group (n = 20 rats total) to quantify protein levels. Tissue from 1 aged SD-fed rat, however, was unrecoverable and not included in the analysis. The membrane and soluble fractions were isolated and quantified according to previously published procedures (Hernandez et al., 2018a, 2018b, 2018c; McQuail et al., 2012). ~5 μg of protein per lane was separated on 4-15% TGX gels (Bio-Rad Laboratories, Hercules, CA, USA) at 200V for 40 minutes in tris-glycine running buffer (Bio-Rad). Total protein was transferred to a 0.45 μm pore nitrocellulose membrane at 20 V for 7 minutes using iBlot Gel Transfer Nitrocellulose Stacks (NR13046-01, Invitrogen, Waltham, MA, USA) and an iBlot machine (Invitrogen, Waltham, MA, USA). All experiments were conducted in triplicate, and the loading order of samples was counterbalanced between gels and experiments to control for systematic variation in the electrophoresis and electroblotting procedures.

Immediately after transfer, membranes were stained for total protein using LiCor’s Revert total protein stain for 5 minutes (Li-Cor, 926-11011) and scanned using a 685 nm laser on an Odyssey IR Scanner (Li-Cor, Lincoln, Nebraska USA) to detect total protein levels. Membranes were then placed into Rockland blocking buffer (Rockland Blocking Buffer, Rockland, Gilbertsville, PA, USA) for 1 hour at room temperature. After blocking, membranes were incubated at 4°C overnight with antibodies raised against ROCK2 (diluted 1:10,000, abcam #ab125025). Membranes were washed in tris buffered saline before incubation in donkey anti-rabbit secondary antibodies conjugated to IRDye800 (diluted 1:15000; LI-COR). Blots were scanned using a 785 nm laser on an Odyssey IR Scanner. The target band signal to total protein signal ratio was calculated for each technical replicate and data from each independent biological sample were transformed to percent expression of the young, SD-fed group (that is, mean of this group is set to 100%) for statistical analysis.

### Statistical analysis

All data are expressed as the group median with interquartile ranges. Differences across age and diet groups were analyzed using a two-factor ANOVA with the between-subjects factors of age (2 levels: young and aged) and diet (2 levels: standard diet and ketogenic diet). When applicable, hippocampal subregions were analyzed independently. The null hypothesis was rejected at the level of p > 0.05 unless it was Bonferroni corrected for multiple comparisons. All analyses were performed with the Statistical Package for the Social Sciences v25 (IBM, Armonk, NY) or GraphPad Prism version 7.03 for Windows (GraphPad Software, La Jolla, California USA).

## Results

### Confirmation of nutritional ketosis

At the time of sacrifice and tissue collection, all rats on the ketogenic diet (KD) had significantly higher levels of β-hydroxybutyrate (BHB), the primary ketone body in the blood, relative to standard diet-fed (SD) rats (F_[1,26]_ = 89.84; p < 0.0001; Figure 1A). Additionally, all rats on the ketogenic diet had significantly lower levels of circulating glucose relative to SD-fed rats (F_[1,26]_ = 12.55; p = 0.002; Figure 1B). Neither BHB nor glucose significantly differed across age groups at this 12-week time point (F_[1,26]_ = 1.41; p = 0.25 and F_[1,26]_ = 0.54; p = 0.47, respectively). The ratio of glucose to ketone body levels can provide a single value quantification of inferred level of ketosis, which has been shown to correlate with cancer treatment efficacy (Meidenbauer et al., 2015). Therefore, the glucose ketone index (GKI) was also calculated as glucose (mmol/L) / BHB (mml/L). Thus, a lower GKI indicates higher levels of ketosis. The GKI was significantly lower in all KD-fed rats relative to SD-fed rats, regardless of age (F_[1,26]_ = 59.91; p < 0.0001; Figure 1C). After 12 weeks on the diet, there was no significant effect of age on GKI (F_[1,26]_ = 0.45; p = 0.51), and age and diet groups did not interact significantly for any of the three variables (p ≥ 0.40 for all).

**Figure 1:**
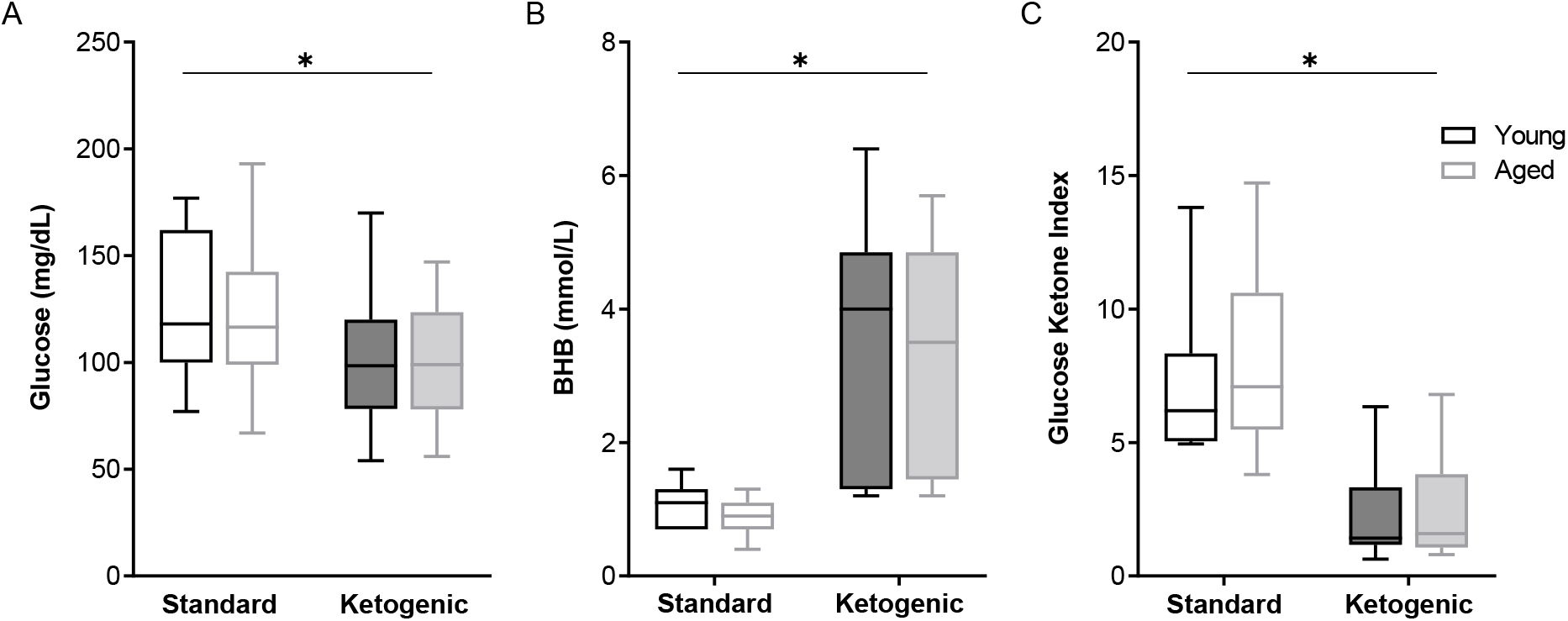
The ketogenic diet-fed rats had significantly higher (A) BHB, lower (B) glucose and lower (C) GKI relative to standard diet-fed rats, regardless of age group. Boxes represent the upper and lower quartiles around the median, and the whiskers are the minimum and maximum observed values.

### Effects of age and diet on hippocampal subregion gene transcript levels

A two-factor ANOVA (age × diet) confirmed that the house keeping gene *RPLP1* did not differ by age or diet group across any region (largest mean difference was <0.425 C_t_, Figure 2; main effects of age: F-values_[1,20-23]_ = 0.891-1.762, p-values = 0.198-0.357; main effects of diet: F-values_[1,20-23]_ = 0.014-1.927, p-values = 0.178-0.907; age × diet interactions: F-values_[1,20-23]_ = 1.252-3.669, p-values = 0.069-0.275).

**Figure 2:**
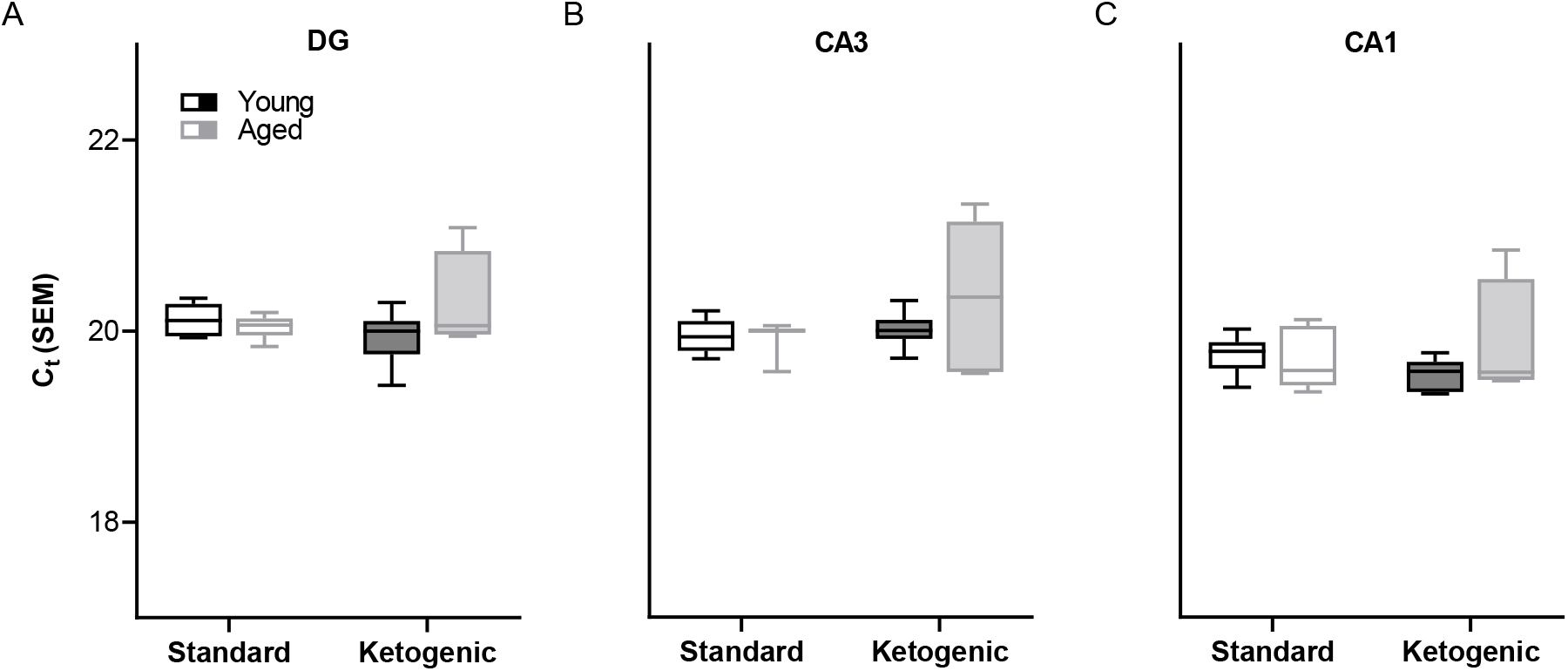
Cycle threshold for housekeeping gene *RPLP1*. Shows the cycle threshold (C_t_) across age and diet groups for the housekeeping gene *RPLP1* in all hippocampal subregions examined. In the dentate gyrus (DG), CA3 and CA1 subregions, the C_t_ did not vary across age and diet groups, nor was there a significant interaction. Thus, *RPLP1* was utilized as a housekeeping gene to normalize the quantification of RNA levels for all other genes of interest.

In the dentate gyrus (DG), a two-factor ANOVA (age × diet) detected that 40 gene transcripts were significantly reduced in the KD group compared to animals on the SD (F-values_[1,21-22]_ = 11.418-22.921, p-values_[FDR-adjusted]_ < 0.005; Figure 3). Figure 3 shows the relative expression levels of these 40 genes for the different age and diet groups. Interestingly, the main effects of age did not reach statistical significance for any of the genes examined (F-values_[1,21-22]_ = 0.001-6.881, p-values_[FDR-adjusted]_ > 0.005), nor were any significant age × diet interactions detected (F-values_[1,21-22]_ = 0.010-3.934, p-values_[FDR-adjusted]_ > 0.005). Analysis of the genes that were significantly affected by diet using the database for annotation, visualization and integrated discovery (DAVID; Huang et al., 2009b, 2009a) identified that 21 of the genes affected by diet contributed to the top ranking and reliable enrichment of the glutamatergic synapse pathway (p_(FDR-adjusted)_=6.632×10^−25^; Enrichr validation: p_(FDR-adjusted)_=1.291×10^−35^; Figure D). We further categorized the genes contributing to the enrichment of this pathway into presynaptic glutamate regulation, postsynaptic excitation, and post-synaptic plasticity. Figure 4 shows the z-scored values for these genes by category across the different diet and age groups. Other pathways most significantly associated with genes affected by the KD in the DG are presented in Table 2.

**Figure 3:**
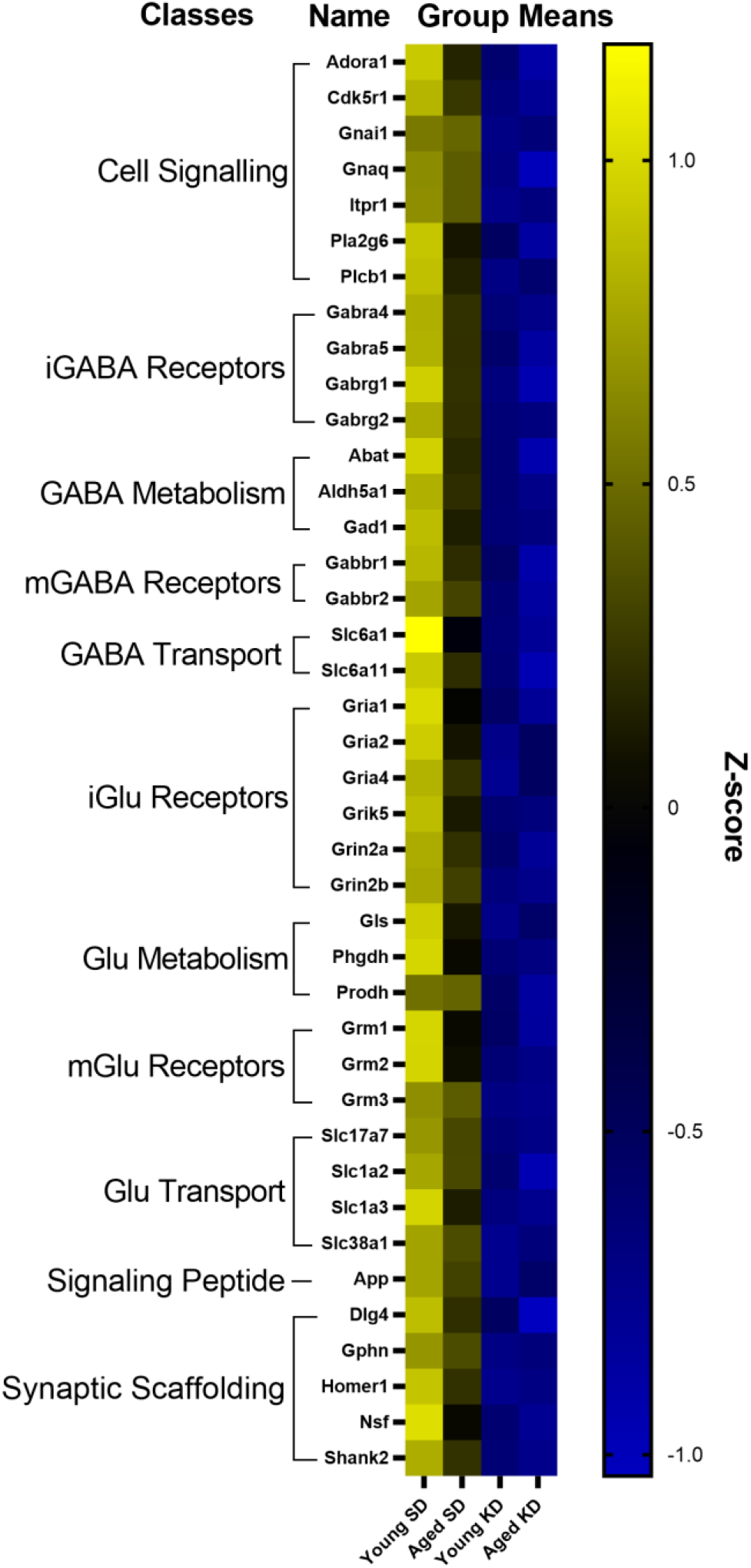
Genes affected by diet in the dentate gyrus. The heat map represents mean z-scores of all genes that were significantly reduced (FDR q = 0.005 threshold) as a function of diet. Genes were categorized into biologically-relevant processes, and experimental groups were ordered to reflect the main effect of diet. Yellow represents greater gene expression, whereas blue represents lower gene expression. Black represents no change.

**Figure 4:**
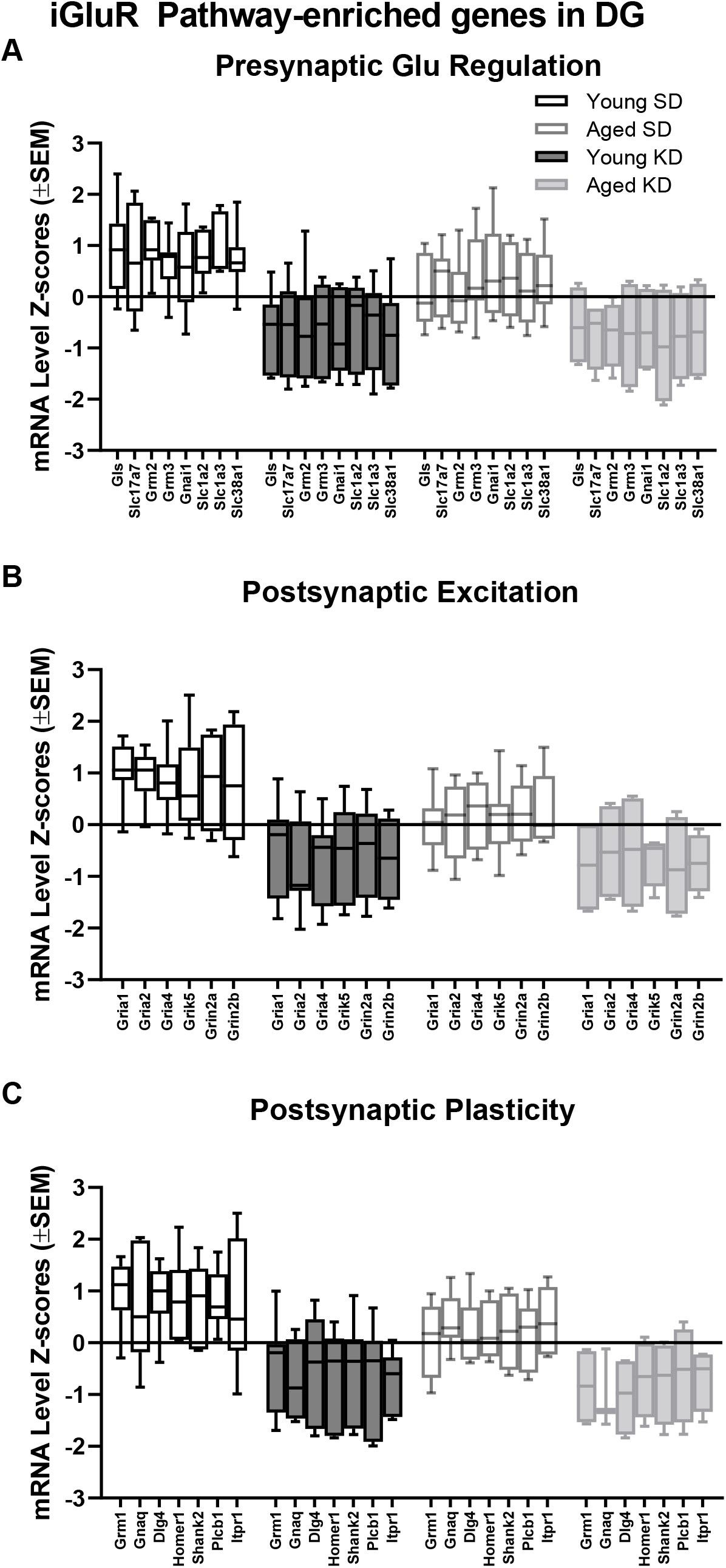
Genes reliably changed with diet that contribute to dentate gyrus ionotropic glutamate receptor (glutamatergic synapse) pathway enrichment. A) 8 genes associated with presynaptic glutamate regulation were reduced as a function of diet. B) 6 genes associated with postsynaptic excitation were reduced as a function of diet. C) 7 genes associated with postsynaptic plasticity were reduced as a function of diet. Black bars represent young; gray bars represent aged; open bars represent standard diet; and filled bars represent ketogenic diet. Error bars represent the interquartile range (IQR).

**Table 2:** Pathways significantly associated with the gene expression changes observed within the dentate gyrus of rats on a ketogenic diet as identified by DAVID.

| TERM | GENE COUNT | %GENES CONTRIBUTING | GENES CONTRIBUTING TO ENRICHMENT | P VALUE | FDR-ADJUSTED P-VALUE | FOLD ENRICHMENT |
| --- | --- | --- | --- | --- | --- | --- |
| :GLUTAMATERGIC SYNAPSE | 21 | 53.85 | GNAI1, GRIK5, GRIN2A, GRIA4, HOMER1, SHANK2, GRM1, ITPR1, SLC17A7, SLC1A2, GRM3, SLC1A3, GRM2, GNAQ, GRIA2, GRIN2B, GRIA1, GLS, DLG4, SLC38A1, PLCB1 | 6.14 E-28 | 6.63 E-25 | 37.24 |
| GABAERGIC SYNAPSE | 14 | 35.90 | GABRG1, GABRG2, SLC6A1, GABRA4, GNAI1, GABRA5, GABBR1, GABBR2, GPHN, GLS, ABAT, SLC38A1, GAD1, NSF | 5.01 E-17 | 5.41 E-14 | 32.81 |
| RETROGRADE ENDOCANNABINOID SIGNALING | 13 | 33.33 | GABRG1, GABRG2, GABRA4, GNAI1, GABRA5, GRIA4, GRM1, ITPR1, SLC17A7, GNAQ, GRIA2, GRIA1, PLCB1 | 2.23 E-14 | 2.41 E-11 | 25.74 |
| NICOTINE ADDICTION | 10 | 25.64 | SLC17A7, GABRG1, GABRG2, GRIN2B, GRIA2, GABRA4, GRIA1, GABRA5, GRIN2A, GRIA4 | 1.11 E-13 | 1.20 E-10 | 50.98 |
| NEUROACTIVE LIGAND-RECEPTOR INTERACTION | 16 | 41.03 | GABRG1, GABRG2, GABRA4, GABRA5, GRIK5, GABBR1, GRIN2A, GABBR2, GRIA4, ADORA1, GRM1, GRM3, GRM2, GRIA2, GRIN2B, GRIA1 | 1.36 E-12 | 1.47 E-09 | 11.17 |
| COCAINE ADDICTION | 8 | 20.51 | GRM3, CDK5R1, GRM2, GRIN2B, GRIA2, GNAI1, GRIN2A, DLG4 | 1.45 E-09 | 1.57 E-06 | 35.46 |
| CIRCADIAN ENTRAINMENT | 9 | 23.08 | GNAQ, GRIN2B, GRIA2, GNAI1, GRIA1, GRIN2A, GRIA4, PLCB1, ITPR1 | 1.29 E-08 | 1.39 E-05 | 18.92 |
| LONG-TERM POTENTIATION | 8 | 20.51 | GNAQ, GRIN2B, GRIA2, GRIA1, GRIN2A, PLCB1, GRM1, ITPR1 | 1.77 E-08 | 1.92 E-05 | 25.10 |
| DOPAMINERGIC SYNAPSE | 9 | 23.08 | GNAQ, GRIN2B, GRIA2, GNAI1, GRIA1, GRIN2A, GRIA4, PLCB1, ITPR1 | 1.22 E-07 | 1.32 E-04 | 14.23 |
| MORPHINE ADDICTION | 8 | 20.51 | GABRG1, GABRG2, GABRA4, GNAI1, GABRA5, GABBR1, GABBR2, ADORA1 | 2.03 E-07 | 2.20 E-04 | 17.73 |
| LONG-TERM DEPRESSION | 7 | 17.95 | GNAQ, GRIA2, GNAI1, GRIA1, PLCB1, GRM1, ITPR1 | 3.93 E-07 | 4.24 E-04 | 23.02 |
| CAMP SIGNALING PATHWAY | 9 | 23.08 | GRIN2B, GRIA2, GNAI1, GRIA1, GABBR1, GRIN2A, GRIA4, GABBR2, ADORA1 | 2.88 E-06 | 0.00 | 9.41 |
| ESTROGEN SIGNALING PATHWAY | 7 | 17.95 | GNAQ, GNAI1, GABBR1, GABBR2, PLCB1, GRM1, ITPR1 | 5.27 E-06 | 0.01 | 14.87 |
| AMYOTROPHIC LATERAL SCLEROSIS (ALS) | 5 | 12.82 | SLC1A2, GRIN2B, GRIA2, GRIA1, GRIN2A | 1.26 E-04 | 0.14 | 18.54 |
| ALZHEIMER'S DISEASE | 7 | 17.95 | CDK5R1, APP, GNAQ, GRIN2B, GRIN2A, PLCB1, ITPR1 | 1.96 E-04 | 0.21 | 7.84 |
| AMPHETAMINE ADDICTION | 5 | 12.82 | GRIN2B, GRIA2, GRIA1, GRIN2A, GRIA4 | 2.28 E-04 | 0.25 | 15.93 |
| RENIN SECRETION | 5 | 12.82 | GNAQ, GNAI1, PLCB1, ADORA1, ITPR1 | 2.72 E-04 | 0.29 | 15.22 |
| ALANINE, ASPARTATE AND GLUTAMATE METABOLISM | 4 | 10.26 | ALDH5A1, GLS, ABAT, GAD1 | 5.90 E-04 | 0.64 | 23.31 |
| GAP JUNCTION | 5 | 12.82 | GNAQ, GNAI1, PLCB1, GRM1, ITPR1 | 7.70 E-04 | 0.83 | 11.59 |
| SEROTONERGIC SYNAPSE | 5 | 12.82 | APP, GNAQ, GNAI1, PLCB1, ITPR1 | 0.00 | 2.92 | 8.22 |
| GASTRIC ACID SECRETION | 4 | 10.26 | GNAQ, GNAI1, PLCB1, ITPR1 | 0.00 | 5.23 | 11.17 |
| CGMP-PKG SIGNALING PATHWAY | 5 | 12.82 | GNAQ, GNAI1, PLCB1, ADORA1, ITPR1 | 0.01 | 7.42 | 6.29 |
| BUTANOATE METABOLISM | 3 | 7.69 | ALDH5A1, ABAT, GAD1 | 0.01 | 8.07 | 21.85 |
| CALCIUM SIGNALING PATHWAY | 5 | 12.82 | GNAQ, GRIN2A, PLCB1, GRM1, ITPR1 | 0.01 | 11.49 | 5.51 |
| HUNTINGTON'S DISEASE | 5 | 12.82 | GNAQ, GRIN2B, DLG4, PLCB1, ITPR1 | 0.02 | 15.41 | 5.02 |
| CHOLINERGIC SYNAPSE | 4 | 10.26 | GNAQ, GNAI1, PLCB1, ITPR1 | 0.02 | 15.98 | 7.28 |
| INFLAMMATORY MEDIATOR REGULATION OF TRP CHANNELS | 4 | 10.26 | GNAQ, PLA2G6, PLCB1, ITPR1 | 0.02 | 17.05 | 7.09 |
| RAP1 SIGNALING PATHWAY | 5 | 12.82 | GNAQ, GRIN2B, GNAI1, GRIN2A, PLCB1 | 0.02 | 18.62 | 4.72 |
| VASCULAR SMOOTH MUSCLE CONTRACTION | 4 | 10.26 | GNAQ, PLA2G6, PLCB1, ITPR1 | 0.02 | 19.66 | 6.69 |
| SPHINGOLIPID SIGNALING PATHWAY | 4 | 10.26 | GNAQ, GNAI1, PLCB1, ADORA1 | 0.02 | 20.43 | 6.58 |
| PLATELET ACTIVATION | 4 | 10.26 | GNAQ, GNAI1, PLCB1, ITPR1 | 0.03 | 24.86 | 6.04 |
| OXYTOCIN SIGNALING PATHWAY | 4 | 10.26 | GNAQ, GNAI1, PLCB1, ITPR1 | 0.04 | 32.23 | 5.37 |
| THYROID HORMONE SYNTHESIS | 3 | 7.69 | GNAQ, PLCB1, ITPR1 | 0.04 | 36.75 | 9.00 |
| SALIVARY SECRETION | 3 | 7.69 | GNAQ, PLCB1, ITPR1 | 0.05 | 43.01 | 8.05 |
| ALDOSTERONE SYNTHESIS AND SECRETION | 3 | 7.69 | GNAQ, PLCB1, ITPR1 | 0.06 | 49.10 | 7.28 |
| GNRH SIGNALING PATHWAY | 3 | 7.69 | GNAQ, PLCB1, ITPR1 | 0.07 | 54.91 | 6.65 |
| PANCREATIC SECRETION | 3 | 7.69 | GNAQ, PLCB1, ITPR1 | 0.08 | 57.68 | 6.37 |
| MELANOGENESIS | 3 | 7.69 | GNAQ, GNAI1, PLCB1 | 0.08 | 59.70 | 6.18 |
| GLUCAGON SIGNALING PATHWAY | 3 | 7.69 | GNAQ, PLCB1, ITPR1 | 0.08 | 60.36 | 6.12 |
| CHAGAS DISEASE (AMERICAN TRYPANOSOMIASIS) | 3 | 7.69 | GNAQ, GNAI1, PLCB1 | 0.09 | 64.81 | 5.72 |

In contrast to DG, a two-factor ANOVA (age × diet) did not detect a significant main effect of diet on the expression of synapse-related genes in CA3 (F-values_[1,20-23]_ = 1.259-15.734, p-values_(FDR-adjusted)_ > 0.005). Age group, however, did significantly affect 24 gene transcripts, with aged rats having significantly lower expression levels compared to young animals (F-values_[1,20-23]_ = 12.683-28.988, p-values_[FDR-adjusted]_ < 0.005; Figure 5). The interaction effect of diet and age group did not reach statistical significance for any of the genes examined in CA3 (F-values_[1,20-23]_ = 0.001-4.028, p-values_(FDR-adjusted)_ > 0.005). Analysis of genes affected by age using DAVID identified that 12 of these genes contributed to the top ranking and reliable enrichment of the glutamatergic synapse pathway (p_(FDR-adjusted)_=2.975×10^−12^; Enrichr validation: p_(FDR-adjusted)_=5.055×10^−26^; Figure 5). Genes contributing to the enrichment of this pathway were categorized into presynaptic glutamate regulation, postsynaptic excitation, and post-synaptic plasticity (Figure 6). Other pathways most significantly associated with these genes are presented in Table 3.

**FIGURE 5:**
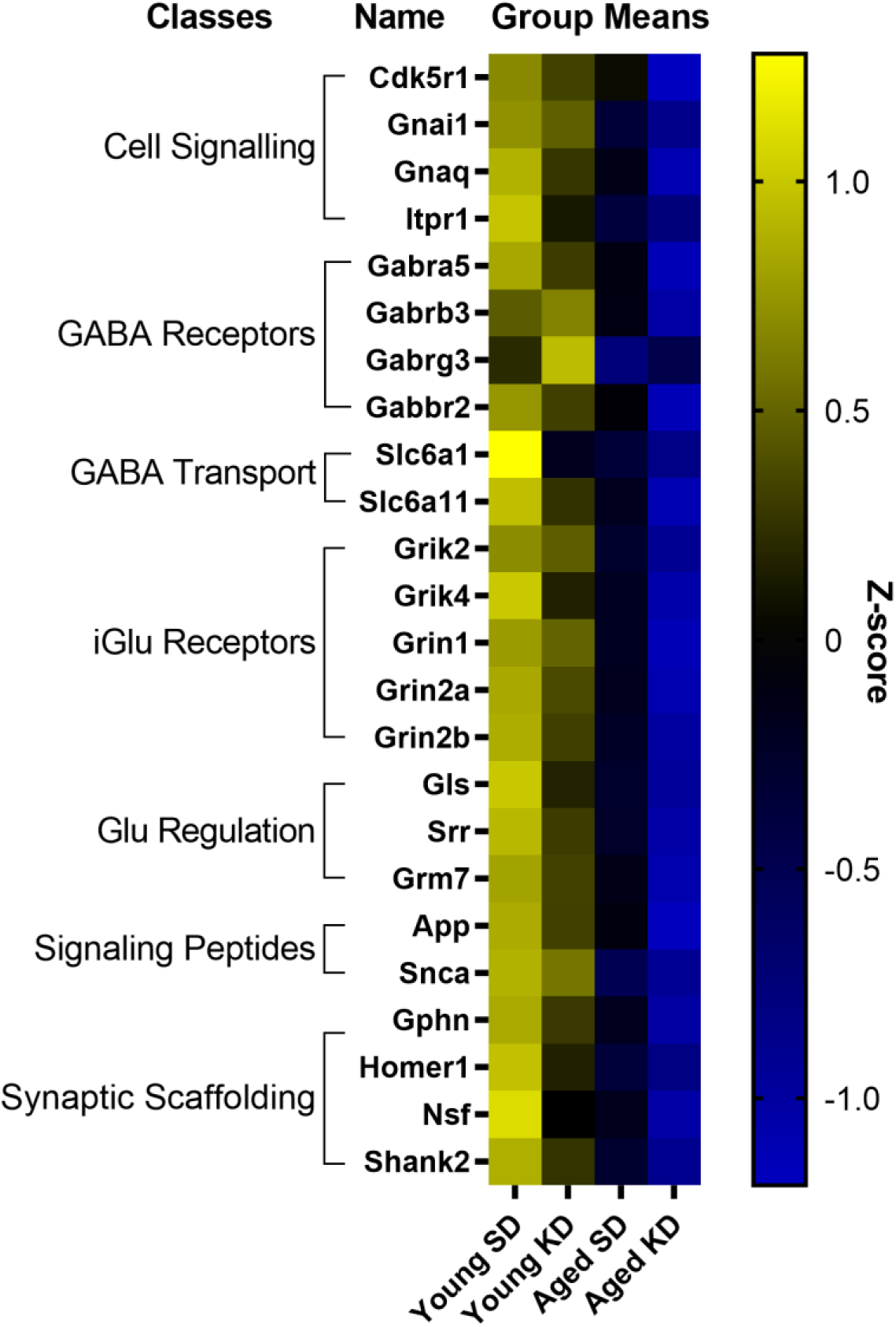
Genes affected by age in CA3. The heat map represents mean z-scores of all genes that were significantly reduced (FDR q = 0.005 threshold) as a function of age. Genes were categorized into biologically-relevant processes, and experimental groups were ordered to reflect the main effect of age. Yellow represents greater gene expression, whereas blue represents lower gene expression. Black represents no change.

**FIGURE 6:**
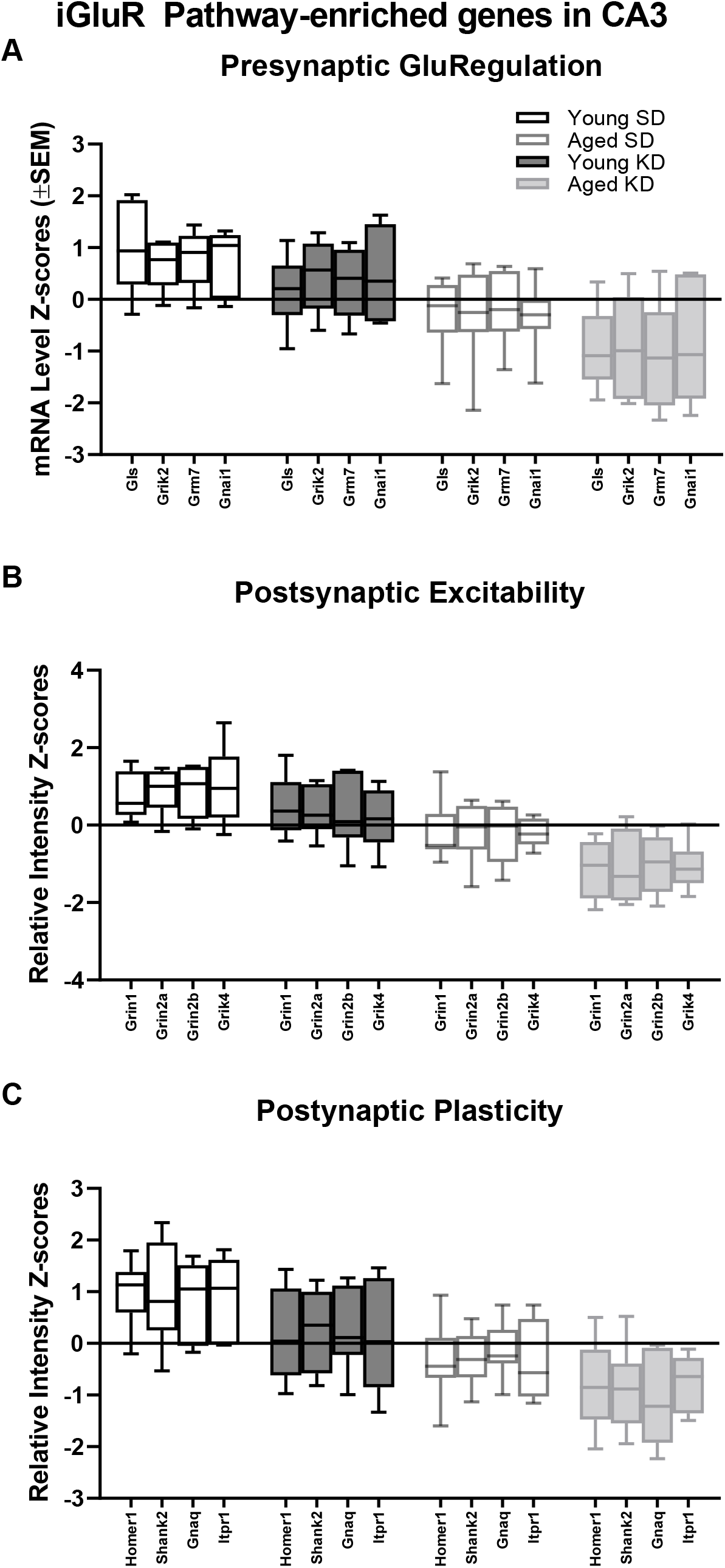
Genes reliably changed with age that contribute to CA3 ionotropic glutamate receptor (glutamatergic synapse) pathway enrichment. A) 4 genes associated with presynaptic glutamate regulation were reduced as a function of age. B) 4 genes associated with postsynaptic excitation were reduced as a function of age. C) 4 genes associated with postsynaptic plasticity were reduced as a function of age. Black bars represent young; gray bars represent aged; open bars represent standard diet; and filled bars represent ketogenic diet. Error bars represent the interquartile range (IQR).

**Table 3:** Pathways significantly associated with the gene expression changes observed within CA3 of aged rats as identified by DAVID.

| TERM | GENE COUNT | %GENES CONTRIBUTING | GENES CONTRIBUTING TO ENRICHMENT | P VALUE | FDR-ADJUSTED P-VALUE | FOLD ENRICHMENT |
| --- | --- | --- | --- | --- | --- | --- |
| GLUTAMATERGIC SYNAPSE | 12 | 0.31 | GNAQ, GRIN2B, GNAI1, GRIK2, GRM7, GLS, GRIK4, GRIN1, GRIN2A, HOMER1, SHANK2, ITPR1 | 2.87 E-15 | 2.98 E-12 | 35.16 |
| GABAERGIC SYNAPSE | 9 | 0.23 | GPHN, GABRG3, GABRB3, SLC6A1, GNAI1, GLS, GABRA5, GABBR2, NSF | 5.12 E-11 | 5.28 E-08 | 34.85 |
| NEUROACTIVE LIGAND-RECEPTOR INTERACTION | 10 | 0.26 | GABRG3, GABRB3, GRIN2B, GRIK2, GRM7, GRIK4, GRIN1, GABRA5, GRIN2A, GABBR2 | 4.38 E-08 | 4.52 E-05 | 11.54 |
| NICOTINE ADDICTION | 6 | 0.15 | GABRG3, GABRB3, GRIN2B, GRIN1, GABRA5, GRIN2A | 6.99 E-08 | 7.21 E-05 | 50.54 |
| ALZHEIMER'S DISEASE | 8 | 0.21 | CDK5R1, APP, GNAQ, GRIN2B, GRIN1, SNCA, GRIN2A, ITPR1 | 4.45 E-07 | 4.59 E-04 | 14.81 |
| CIRCADIAN ENTRAINMENT | 6 | 0.15 | GNAQ, GRIN2B, GNAI1, GRIN1, GRIN2A, ITPR1 | 6.16 E-06 | 0.01 | 20.84 |
| COCAINE ADDICTION | 5 | 0.13 | CDK5R1, GRIN2B, GNAI1, GRIN1, GRIN2A | 7.35 E-06 | 0.01 | 36.62 |
| RETROGRADE ENDOCANNABINOID SIGNALING | 6 | 0.15 | GABRG3, GNAQ, GABRB3, GNAI1, GABRA5, ITPR1 | 8.28 E-06 | 0.01 | 19.63 |
| LONG-TERM POTENTIATION | 5 | 0.13 | GNAQ, GRIN2B, GRIN1, GRIN2A, ITPR1 | 2.94 E-05 | 0.03 | 25.92 |
| MORPHINE ADDICTION | 5 | 0.13 | GABRG3, GABRB3, GNAI1, GABRA5, GABBR2 | 1.16 E-04 | 0.12 | 18.31 |
| SEROTONERGIC SYNAPSE | 5 | 0.13 | APP, GNAQ, GABRB3, GNAI1, ITPR1 | 3.66 E-04 | 0.38 | 13.59 |
| DOPAMINERGIC SYNAPSE | 5 | 0.13 | GNAQ, GRIN2B, GNAI1, GRIN2A, ITPR1 | 4.25 E-04 | 0.44 | 13.06 |
| CAMP SIGNALING PATHWAY | 5 | 0.13 | GRIN2B, GNAI1, GRIN1, GRIN2A, GABBR2 | 0.00 | 2.04 | 8.64 |
| ESTROGEN SIGNALING PATHWAY | 4 | 0.10 | GNAQ, GNAI1, GABBR2, ITPR1 | 0.00 | 2.44 | 14.04 |
| RAP1 SIGNALING PATHWAY | 5 | 0.13 | GNAQ, GRIN2B, GNAI1, GRIN1, GRIN2A | 0.00 | 2.95 | 7.80 |
| AMYOTROPHIC LATERAL SCLEROSIS (ALS) | 3 | 0.08 | GRIN2B, GRIN1, GRIN2A | 0.01 | 10.26 | 18.38 |
| ALCOHOLISM | 4 | 0.10 | GRIN2B, GNAI1, GRIN1, GRIN2A | 0.01 | 12.55 | 7.66 |
| LONG-TERM DEPRESSION | 3 | 0.08 | GNAQ, GNAI1, ITPR1 | 0.01 | 12.75 | 16.30 |
| AMPHETAMINE ADDICTION | 3 | 0.08 | GRIN2B, GRIN1, GRIN2A | 0.01 | 13.49 | 15.79 |
| CALCIUM SIGNALING PATHWAY | 4 | 0.10 | GNAQ, GRIN1, GRIN2A, ITPR1 | 0.01 | 14.23 | 7.28 |
| RENIN SECRETION | 3 | 0.08 | GNAQ, GNAI1, ITPR1 | 0.02 | 14.64 | 15.09 |
| GASTRIC ACID SECRETION | 3 | 0.08 | GNAQ, GNAI1, ITPR1 | 0.02 | 17.01 | 13.85 |
| HUNTINGTON'S DISEASE | 4 | 0.10 | GNAQ, GRIN2B, GRIN1, ITPR1 | 0.02 | 17.87 | 6.64 |
| GAP JUNCTION | 3 | 0.08 | GNAQ, GNAI1, ITPR1 | 0.03 | 23.33 | 11.49 |
| CHOLINERGIC SYNAPSE | 3 | 0.08 | GNAQ, GNAI1, ITPR1 | 0.04 | 34.11 | 9.02 |
| PLATELET ACTIVATION | 3 | 0.08 | GNAQ, GNAI1, ITPR1 | 0.06 | 44.50 | 7.49 |
| OXYTOCIN SIGNALING PATHWAY | 3 | 0.08 | GNAQ, GNAI1, ITPR1 | 0.07 | 51.84 | 6.65 |
| CGMP-PKG SIGNALING PATHWAY | 3 | 0.08 | GNAQ, GNAI1, ITPR1 | 0.08 | 55.96 | 6.24 |

Finally, within CA1, there were no significant effects of age or diet on gene transcript levels, nor were there any significant age × diet interactions after adjusting for a FDR of q = 0.005 (two-factor, age × diet ANOVA: F-values_[1,18-20]_ = 0.000-8.629, ps_(FDR-adjusted)_>0.005).

### The ketogenic diet eestores age-related increases in hippocampal ROCK2 expression

The reduction in gene transcripts associated with glutamatergic signaling in the DG suggests that long-term nutritional ketosis could decrease excitatory synaptic transmission. Previous work examining protein levels of the vesicular glutamate transporter 1 in whole hippocampal homogenates, however, showed that the KD remediated age-related loss of this essential synaptic protein without otherwise influencing expression in young rats (Hernandez et al., 2018a). This previous work suggests that there is not an overall loss of excitatory synapses within the hippocampus following the KD, and that there may even be a restoration of synapse number in aged rats. In the hippocampus, most excitatory synapses are on dendritic spines (McKinney, 2010). Rho-associated coiled-coil containing protein kinases (ROCKs) regulate the actin cytoskeleton at these synapses, and inhibition of these molecules increases spine density (Swanger et al., 2016). These data suggest that elevated ROCK levels in the hippocampus could potentially be related to excitatory synapse loss. Furthermore, inhibition of ROCK improves spatial working memory in aged rats suggesting that decreasing ROCK may be beneficial for the treatment of cognitive aging or neurodegenerative disorders resulting in dendritic spine alterations (Henderson et al., 2016; Kubo et al., 2008). Therefore, ROCK2 protein levels within the soluble fraction of hippocampal homogenates were quantified via Western Blotting. Within the hippocampus, there was a significant age-dependent increase in ROCK2 expression in aged rats on the SD relative to young animals on the SD (F_[1,7]_ = 11.61; p = 0.01; Figure 7). Because of the main effect of age across SD-fed subjects, the possibility that a KD could restore these levels was explored using a multivariate ANOVA. While there were no main effects of either age (F_[1,15]_ = 2.21; p = 0.16) or diet (F_[1,15]_ = 4.36; p = 0.054), there was a significant interaction between age and diet (F_[1,15]_ = 11.74; p = 0.004). Furthermore, when young and aged rats on KD were directly compared, there was no longer an effect of age (F_[1,8]_ = 1.97; p = 0.20). These data show that the KD was able to reduce levels of ROCK2 in the hippocampus of aged animals to levels comparable to those observed in young rats. This observation provides indirect evidence that the KD does not lead to an overall reduction of excitatory synapse number. Thus, the diet related changes within the dentate gyrus are likely due to atered regulation of glutamatergic synapses without a reduction in excitatory synapses. Functional assessment of hippocampal neurophysiology will be necessary to determine the impact of long-term nutritional ketosis on the aged hippocampus.

**Figure 7.**
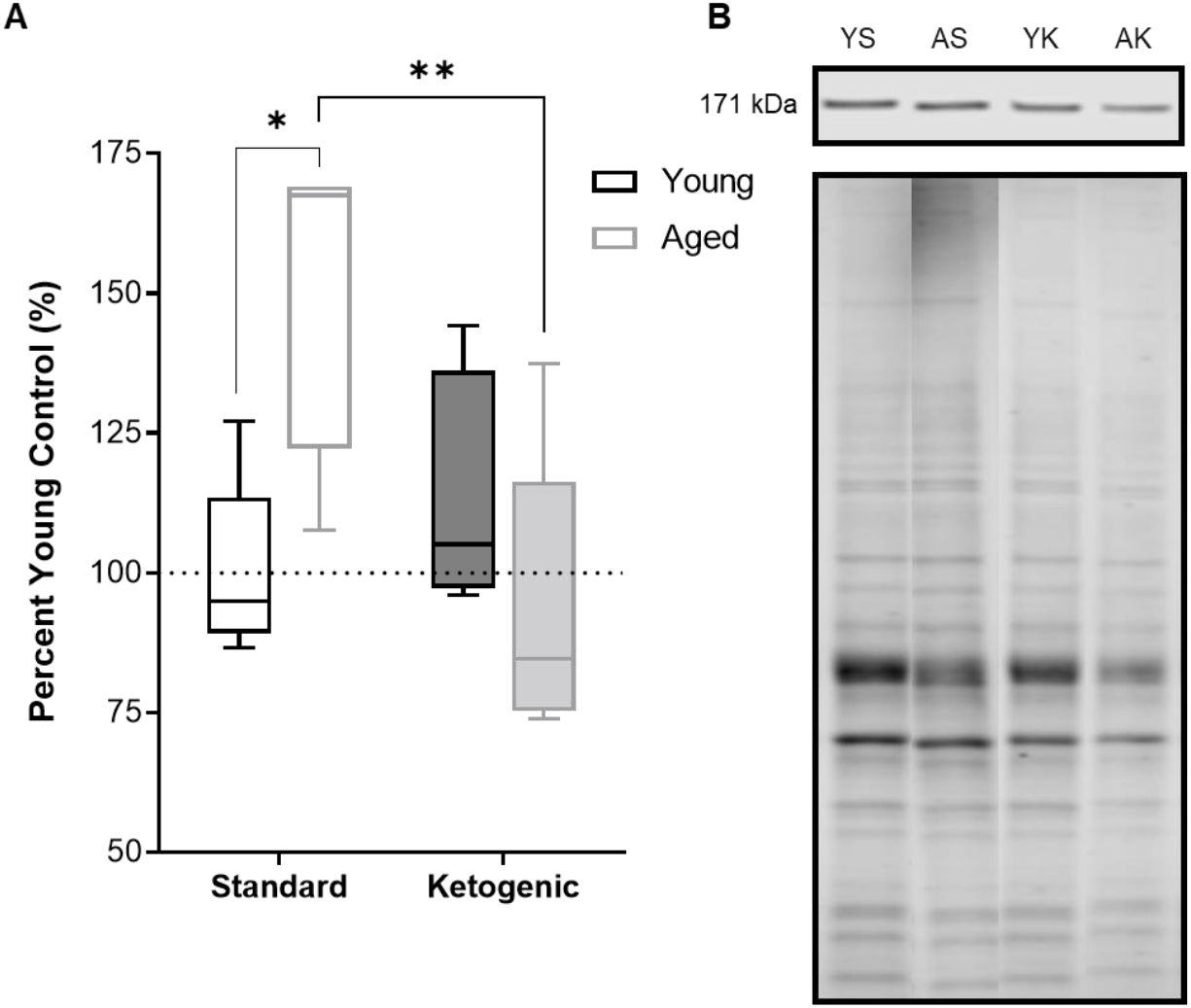
Hippocampal ROCK2 protein expression. (A) Aged rats demonstrated significantly higher levels of ROCK2 expression in the hippocampus relative to young standard-fed (SD) rats, but levels were restored to that of the young SD-fed rats in ketogenic-fed (KD) aged subjects. (B) Representative ROCK2 bands (top), which were then normalized to total protein stains (bottom) to account for variability in total protein loading. Note total protein was less in aged compared to young animals. Boxes represent the interquartile range, and the whiskers are the minimum and maximum observed values. All values are expressed as percent of young SD-fed controls (dotted line).

## Discussion

The goal of this study was to investigate the effects of age and diet on glutamatergic and GABAergic signaling-related gene expression from cell body-enriched tissue samples across subregions of the hippocampus. There was a significant reduction in transcript levels of several genes associated with pre- and post-synaptic signaling mechanisms within the dentate gyrus (DG) of ketogenic diet (KD)-fed rats relative to standard diet (SD)-fed rats irrespective of age. Notably, several gene targets reduced as a function of diet in the DG were also reduced in CA3 of aged rats relative to young rats irrespective of diet. Consistent with previous work, there were no effects of age on transcript levels detected within CA1 (Haberman et al., 2011; Tran et al., 2018; but see Zeier et al., 2011). The current findings build on this to show that CA1 synapse-related transcript levels were also not significantly altered by long-term dietary ketosis. Interestingly, a pathway enrichment analysis highlighted the age-related vulnerability of CA3 glutamatergic synapses in addition to a potential diet-dependent compensation within DG glutamatergic synapses (see Figure 8).

**Figure 8:**
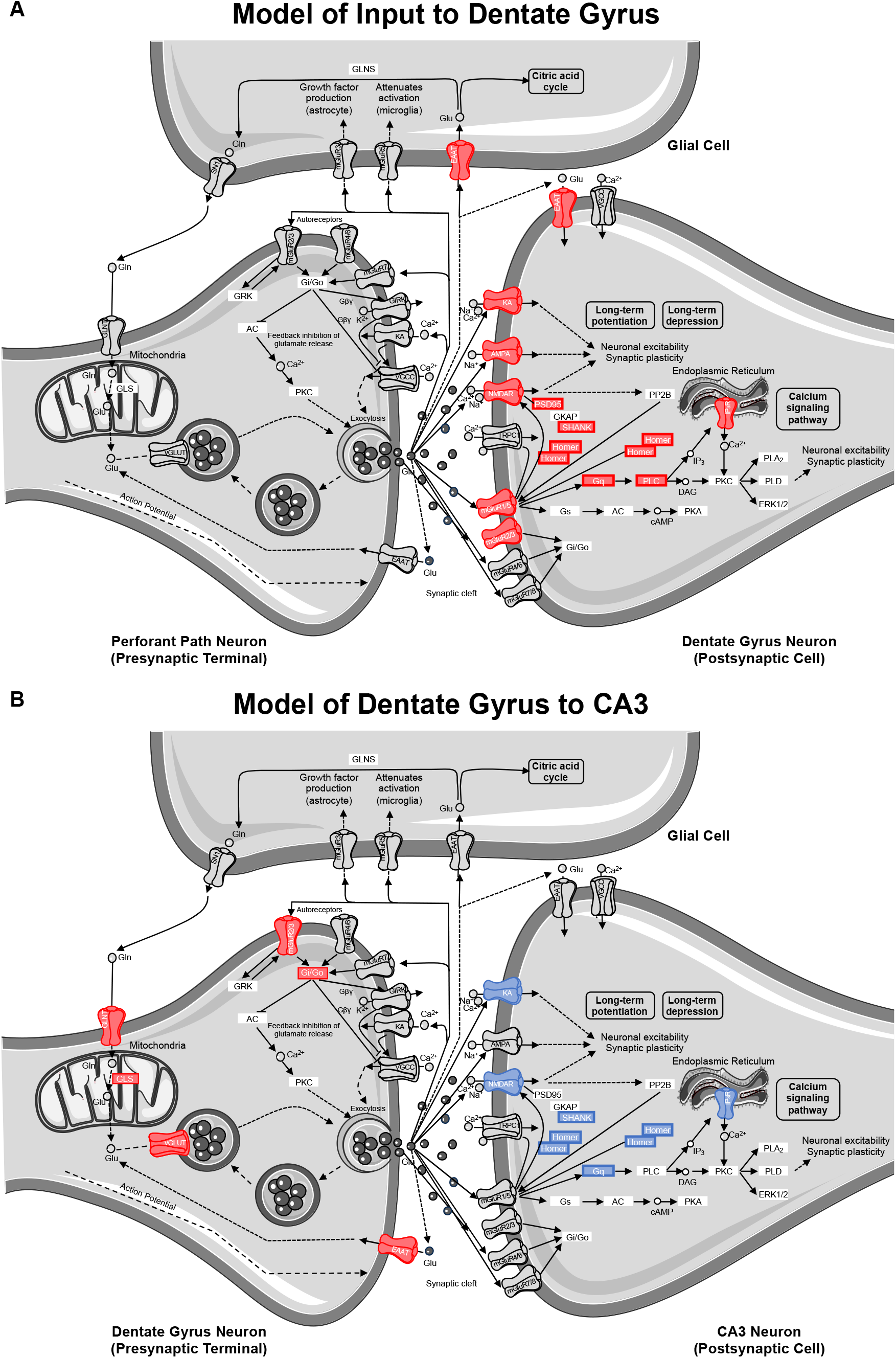
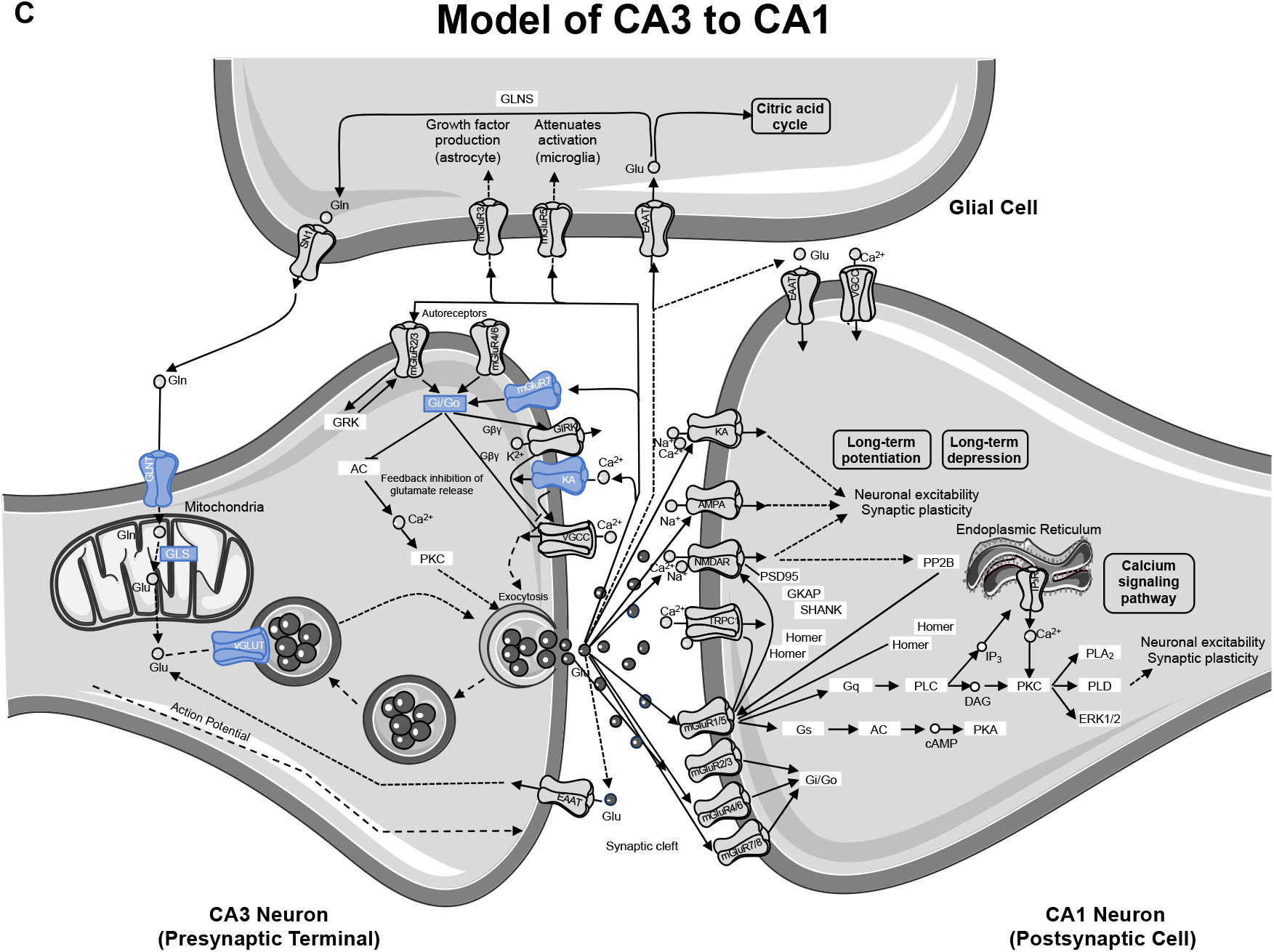
Model of gene contributions to significant ionotropic glutamate receptor (glutamatergic synapse; KEGG) pathway enrichment as determined by DAVID. Significant reductions in receptors/genes by diet (red) and age (blue) are indicated by color.

### Reduction of gene transcript levels by ketogenic diet is specific to dentate gyrus

Although the DG receives input from several cortical and subcortical regions, its primary excitatory input is from the perforant pathway emanating from layer II neurons of the medial and lateral entorhinal cortices and synapsing in the outer two-thirds of the molecular layer (Witter and Amaral, 2004). The inner molecular layer receives commissural and other non-entorhinal cortical inputs, including inhibitory basket cell projections (Sik et al., 1997). The tissue isolated for quantitative RNA analysis in the current study, however, primarily consisted of DG neuron cell bodies with a smaller contribution from hilar cells. Thus, changes in gene expression largely reflect alterations within the DG granule cells densely packed into the principal cell layer, but may also be influenced to some extent by glutamatergic mossy cells (Amaral et al., 2007). Moreover, changes in synapse-related gene transcripts cannot be localized to a specific input layer of the dendritic field that will have variable distributions of inhibitory, excitatory and neuromodulatory input. That being said, transcript reductions in animals on the KD point to potential alterations in granule cell dendritic input (post-synaptic signaling), as well as altered granule cell axonal output from the mossy fibers (presynaptic transmission). Therefore, a pathway enrichment analysis was used to facilitate modeling the most reliable changes to DG dendritic input and DG axonal transmission. According to DAVID’s functional pathway enrichment analyses, the glutamatergic synapse pathway was the most reliably enriched.

Of the 40 genes significantly affected by the KD in the DG, 21 genes contributed to this pathway enrichment. Among these genes, 13 had primarily postsynaptic functions (see Figure 8A), and therefore reflect changes on dendrites receiving input from perforant path axons or other afferents. Specifically, a number of contributors to this pathway enrichment included gene transcripts of postsynaptic proteins integral to neuronal excitability (e.g., ionotropic glutamate receptors), synaptic plasticity (e.g., scaffolding proteins), and second messenger signaling cascades (e.g., metabotropic glutamate receptors and downstream effectors). Together these data suggest that a KD may reduce glutamatergic signaling in granule cell dendrites. This has interesting implications for both aging and epilepsy. In advanced age, the excitatory postsynaptic field potential for a given amplitude of the presynaptic perforant path fiber potential is larger relative to young animals (Barnes and McNaughton, 1980), suggesting that excitatory synapses in aged rats are more powerful. Similarly, intracellular recording methods measuring unitary excitatory postsynaptic potential amplitude in young and aged rats, show that the depolarization resulting from stimulation of a single fiber is increased in old compared to young granule cells (Foster et al., 1991). Thus, it is conceivable that nutritional ketosis may reverse this age-related increase in synaptic strength. Relatedly, the KD is known to be effective at reducing seizure activity (Baranano and Hartman, 2008; Lefevre and Aronson, 2000). Epileptiform activity in medial temporal lobe epilepsy is generated by dentate gyrus granule cells (Bumanglag and Sloviter, 2018). The ketogenic diet may reduce excitatory drive onto granule cells to reset the balance between excitation and inhibition thereby conferring seizure projection. Consistent with the current data showing that the DG is particularly sensitive to a KD, a previous study observed an increased number and size of mitochondria within the DG region of the hippocampus in KD-fed rats. In this study, the authors hypothesized that increasing the quantity of key metabolites involved in oxidative phosphorylation like glutamate and phosphocreatine can enhance ATP production to fuel pumps on the cell membrane. These changes could therefore stabilize the membrane potential and increase resistance to metabolic stress. Using a gene microarray approach, this study also reported an increase in the transcription of genes involved in mitochondrial biogenesis, which the authors concluded might confer anticonvulsant properties within the hippocampus (Bough et al., 2006).

The KD reduced transcript levels of 8 genes that contribute to pathway enrichment of presynaptic proteins integral to glutamate metabolism (e.g., synthesis and packaging) and synaptic glutamate regulation (e.g., autoreceptors and transport) in DG. This observation has implications for how nutritional ketosis may alter mossy fiber axonal transmission to the stratum lucidum of CA3 (see Figure 8B). With age, there are neurophysiological alterations at the mossy fiber-CA3 synapse that include reduced inhibition (Villanueva-Castillo et al., 2017). It is conceivable that when consuming a KD, altered presynaptic glutamate metabolism may decrease excitatory drive to CA3 to ameliorate age-related hyperexcitability (Thomé et al., 2015; Wilson, 2005), but this idea needs to be empirically tested with functional measurements.

While our previous work has shown the diet can restore hippocampal levels of the vesicular glutamate transporter 1 (vGluT1; Hernandez et al., 2018a), the level of transcript encoding vGluT1 (Slc17a7) in the DG was lower in KD-fed relative to SD-fed rats. One possible explanation for the opposing directional changes between protein and transcript levels is that reduced vGluT1 protein was detected from whole-hippocampal homogenates and could have been driven by changes in other subregions. Another possible explanation reconciling contrasting changes between protein and transcript levels is that the KD increased the stability of vGlut1 protein, resulting in a reduced need for gene transcription. Indeed, reductions in levels of a gene transcript are not always translated to a loss of its protein product (Greenbaum et al., 2003; Gygi et al., 1999; Maier et al., 2009). Additional functional assessments of synaptic integrity will be necessary to differentiate between these contrasting possibilities. Importantly, our ROCK2 protein findings suggest a diet-induced increase in synaptic stability, and as such, KD-related reduced transcript levels in the DG may reflect synaptic stability and improved cognitive performance (Hernandez et al., 2018b).

Notably, the results presented here agree with a hippocampal cDNA microarray study on KD-fed rats showing diet-induced reductions in several genes associated with neuronal excitability that overlap with the current study (Noh et al., 2004). Although the null age effects in the DG are consistent with previous work showing CA3-specific changes in gene expression not observed in DG (Haberman et al., 2011), it must be noted that the SD-fed rats in the current study were fed under a time-restricted schedule. This feeding regimen may have improved peripheral metabolism relative to *ad libitum* eating (Mitchell et al., 2016). Previous studies showing age-related changes in the DG were conducted in *ad libitum* fed rats (Ianov et al., 2016), that may have been metabolically compromised due to excessive caloric intake (Martin et al., 2010). The DG is the subregion that is most vulnerable to glucose dysregulation. In fact, individuals with diabetes have reduced basal cerebral blood volume, an indirect measure of oxygen metabolism, in the DG compared to normal study participants. The magnitude of the decline is related to blood glucose levels. In contrast, basal cerebral blood volume in CA1 and CA3 is not affected by diabetes (Wu et al., 2008). It is likely that metabolic changes disproportionately affect the DG due to the high energy demands of adult neurogenesis (Beckervordersandforth, 2017). This cost is further driven by how densely packed the principal cell layer is and the sheer number of neurons within the DG (Amaral et al., 2007).

### Reduction of gene expression with advanced age is specific to CA3

As the CA3 subregion of the hippocampus receives input from the DG, any functional changes within DG likely have direct downstream consequences on CA3. Similar to the DG, the RNA extracted from CA3 was from cell body-enriched samples, and therefore, transcript level changes observed within this subregion predict alterations in dendritic input primarily from DG (post-synaptic signaling) in addition to CA3 axonal output via Schafer collaterals projecting to CA1 (presynaptic transmission). Of the 24 genes that showed a significant reduction in transcript levels in aged compared to young rats, 12 of these genes contributed to glutamatergic synapse pathway enrichment. Specifically, four contribute to presynaptic glutamate regulation, four contribute to postsynaptic signaling transmission, and four contribute to synaptic plasticity (see Figure 8B). Whereas gene transcripts with postsynaptic protein products overrepresented the diet effects in the DG, the effects of age in the CA3 were less discriminate, suggesting broad effects of aging on CA3 gene expression. Indeed, the age-related vulnerability of CA3 neurons is a well-documented phenomenon. The data presented here are consistent with alterations in CA3 excitability observed in humans (Yassa et al., 2010, 2011), monkeys (Thomé et al., 2015), and rats (Maurer et al., 2017; Robitsek et al., 2015; Wilson, 2005). Additionally, changes in gene expression specifically within CA3, and not DG, correlate with cognitive outcomes in aged rats (Haberman et al., 2011).

Interestingly, there were no diet-induced changes in gene expression within CA3 that survived the false discovery rate threshold. Prior work has shown that in whole hippocampal homogenates nutritional ketosis leads to increased levels of protein for the vesicular GABA transporter (vGAT; Hernandez et al., 2018b, 2018a). This led to the hypothesis that the KD’s effectiveness at preventing epileptiform activity (Gasior et al., 2006; Hartman et al., 2007), reducing anxiety-like behavior and improving cognitive function (Hernandez et al., 2018) was through increasing synaptic levels of GABA. Although the gene encoding vGaT (Slc32a1) was not quantified in the present study, diet-induced reductions in transcript levels of genes associated with glutamate synthesis, packaging, and synaptic clearance on presynaptic DG afferents (Figure 8B) suggest that the KD may reduce excitatory afferent drive to CA3, which could confer resilience in aging and epilepsy. This hypothesis needs to be tested with functional studies that directly measure the neurophysiological interactions between the DG and CA3. The complexity of age- and metabolism-related changes demonstrate the need for such studies and require a multifaceted approach beyond the scope of the excitatory and inhibitory gene panel used here.

### Age-related decrease in synaptic stability is restored in ketogenic diet-fed aged rats

In addition to gene expression, whole hippocampal homogenates were used to investigate protein expression of rho-associated coiled-coil containing protein kinase 2 (ROCK2). ROCK2 is an essential regulator of cytoskeletal elements with the ability to modify dendritic spine stability and number, and this protein has been shown to accumulate in the brains of patients with Alzheimer’s disease (Bobo-Jiménez et al., 2017). Excessive ROCK proteins lead to dendritic spine instability, and inhibition of ROCKs restore microtubules at the dendrites, leading to improved dendritic arborization (Bobo-Jiménez et al., 2017; Chen and Firestein, 2007; Li et al., 2000; Nakayama et al., 2000). Furthermore, ROCK inhibition in aged subjects improves cognitive performance on the hippocampal-dependent Morris water maze (Huentelman et al., 2009). We found aged rats fed the SD had significantly increased levels of ROCK2, suggesting decreased stability in axospinous synapses within the hippocampus of aged subjects. Aged rats fed a KD, however, had ROCK2 levels similar to young rats on both diets, demonstrating a restored ability to appropriately regulate dendritic morphology. These data demonstrate a KD, in conjunction with time restricted feeding, exerts changes in neuronal signaling at the levels of both gene and protein expression.

### Conclusions

Although there were no significant changes within CA1, there were alterations in genes encoding presynaptic proteins at CA3 Schafer collateral terminals synapsing onto CA1. Along these lines, the null diet effects on gene expression within CA3 does not mean that this subregion is functionally unaltered. Although the KD did not rescue an age-related decrease in CA3 transcript levels, the KD altered functionally related genes in the DG, which is upstream and sensitive to metabolic perturbations. Consequently, diet-induced decreases in DG transcript levels may be a compensatory mechanism leading to positive change that helps confer resilience in the face of age-related dysfunction in network dynamics between subregions of the hippocampus. This hypothesis needs to be directly tested with *in vivo* approaches to measure DG-CA3 interactions in aging.

Our study also did not detect age-related calcium dyshomeostasis within CA1, as previously described (Bodhinathan et al., 2010; Campbell et al., 1996; Oh et al., 2013; Zeier et al., 2011). We did not see an increase in *ITPR1* gene expression, which is translated into the IP3 receptor protein. Notably, however, the transciprtion of other genes encoding proteins related to calcium regulation that are known to be affected by age were not evaluated in this panel, including voltage-gated L-type calcium channels and ryanodine receptors (Thibault and Landfield, 1996). Alternatively, others have documented age-related shifts in intracellular signaling cascades and their propensity to recruit IP3 receptor-gated calcium pools despite preserved expression of constituent signaling proteins (e.g. Kumar and Foster, 2007, 2014; McQuail et al., 2013; Nicolle et al., 1999).

Finally, differences between this study and previous reports could be due to implementing time-restricted feeding and prevention of overeating, which improves peripheral metabolism (Anton et al., 2018; Golbidi et al., 2017; Manoogian and Panda, 2017), potentially altering neurometabolism in aged rats regardless of diet. The majority of studies utilizing aged rodents have provided access to food *ad libitum* throughout the lifespan, resulting in metabolically impaired subjects (Martin et al., 2010). Thus, free feeding may exacerbate cognitive and neurobiological deficits, increasing the probability of detecting age effects. In fact, it is well documented that excessive energy consumption can lead to cognitive impairments (Bocarsly et al., 2015; Kanoski and Davidson, 2011; Stranahan et al., 2008). Therefore, it is imperative that investigations into the effects of cognitive interventions account for this often overlooked, yet vital component of animal maintenance through the implementation of proper feeding strategies, or at the very least consider rodent metabolic health as a variable in aging studies providing food *ad libitum*.

## Acknowledgements

This research was funded by NIA R01AG060977 (SNB), NIA F31AG058455 (ARH), NIA K01AG061263 (JAM), Pat Tillman Foundation (CMH), McKnight Brain Research Foundation (ARH, CMH, JAM, JLB, SNB), Claude D. Pepper Older Americans Independence Center Scholar Award (P30 AG028740 (JAM, SNB), University of Florida University Scholars Program (LMT, KTC).

